# Posterior parietal cortex guides visual decisions in rats

**DOI:** 10.1101/066639

**Authors:** Angela M. Licata, Matthew T. Kaufman, David Raposo, Michael B. Ryan, John P. Sheppard, Anne K. Churchland

## Abstract

Neurons in putative decision-making structures can reflect both sensory and decision signals, making their causal role in decisions unclear. Here, we tested whether rat posterior parietal cortex (PPC) is causal for processing visual sensory signals or instead for accumulating evidence for decision alternatives. We optogenetically disrupted PPC activity during decision-making and compared effects on decisions guided by auditory vs. visual evidence. Deficits were largely restricted to visual decisions. To further test for visual dominance in PPC, we evaluated electrophysiological responses following individual sensory events and observed much larger responses following visual stimuli than auditory stimuli. Finally, we measured spike count variability during stimulus presentation and decision formation. This sharply decreased, suggesting the network is stabilized by inputs, unlike what would be expected if sensory signals were locally accumulated. Our findings argue that PPC plays a causal role in discriminating visual signals that are accumulated elsewhere.

## Introduction

A large body of work has documented neural responses during perceptual decisions (Roitman and Shadlen, 2002; Churchland et al., 2008; Rishel et al., 2013; Ding, 2015; Hanks et al., 2015). These studies reveal cortical and subcortical structures that might constitute a brain-wide circuit for transforming raw sensory inputs into plans for action. Although transient disruption of activity in these structures could help in assessing their causal role in such a circuit, these types of experiments have been performed rarely and inconclusively. The importance of causal manipulations is underscored by experiments that found no effect of neural disruption on some decisions, even for areas in which neurons reflect decision signals (Suzuki and Gottlieb, 2013; Erlich et al., 2015; Katz et al., 2016).

The role of one candidate area, the posterior parietal cortex (PPC) of rodents, remains particularly ambiguous because existing work paints conflicting pictures of its role in decision-making. Electrophysiological observations demonstrate that PPC is modulated during both auditory (Raposo et al., 2014; Hanks et al., 2015) and visual (Harvey et al., 2012; Raposo et al., 2014) decisions which unfold gradually over about a second. These slow-timecourse signals could reflect evidence accumulation either in PPC or in a remote area with feedback projections to PPC. If evidence accumulation occurs remotely, PPC’s role may instead be to discriminate individual sensory events so that they can be subsequently accumulated over time and used to estimate overall rate. The ability of individual auditory events to drive PPC neurons has been noted as weak (Hanks et al., 2015) but not studied in depth, since recent work has focused on slower modulation over the course of the entire decision. Deficits for visual, but not auditory, decisions observed after PPC inactivation hint at a putative role in discriminating visual events (Raposo et al., 2014). However, these inactivation studies are not entirely conclusive because neural activity was suppressed continuously for 2-3 hours. This leaves open the possibility that a role in evidence accumulation, or a role in detecting auditory events, might have been missed: 2-3 hours of suppression might permit the animal to adjust its strategy, potentially recruiting alternate neural circuits that are not typically involved. Further, existing studies have not fully characterized the nature of the deficits to visual decisions, leaving it unclear whether inactivation affected sensory processing specifically, or instead affected other decision factors.

Here, we examined PPC’s contribution to decision-making by manipulating and more closely measuring neural responses. First, we used a temporally precise optogenetic perturbation method to disrupt neural activity. By disrupting activity during both visual and auditory decisions, we found specific sensory processing effects but little in the way of more general effects on decisions, such as accumulation of evidence or the ability to report choices. A probabilistic decision analysis offered insight into effects of PPC disruption on non-sensory factors that guide decisions, such as a reliance on reward history. Second, we conducted a temporally precise analysis of previously collected electrophysiological data to isolate the impact of individual auditory and visual events on PPC responses, providing an independent and novel assessment of PPC’s role in auditory and visual processing and decision-making. Finally, we leveraged an analysis of trial-to-trial variability that is informative about the underlying computations taking place within an area. All three approaches support the same conclusion: PPC’s contribution to decision circuits is to discriminate visual stimuli so they can be accumulated elsewhere to guide decision-making.

## Results

### Optogenetic disruption of PPC drives deficits in visual decision-making

We optogenetically stimulated PPC neurons expressing channelrhodopsin 2 (Boyden et al., 2005) (ChR2) in all cell types (Figure 1, Figure Supplement 1). This elevates responses of neurons nonspecifically, an approach that is disruptive to the natural activity pattern in areas like PPC in which neurons with heterogeneous response properties are spatially intermixed (Churchland and Shenoy, 2007; Roberts et al., 2012; Rodgers and DeWeese, 2014; Otchy et al., 2015). To probe for a causal role for PPC, we utilized a decision task (Raposo et al., 2012; Sheppard et al., 2013; Raposo et al., 2014) in which freely moving rats judge whether the rate of a 1000 ms series of auditory or visual events is high or low compared to an abstract category boundary (12.5 Hz; Figure 1B). Stimulation took place throughout the 1000 ms series of events. Short-latency changes could be observed in the local field potential (LFP) and the spikes of single, isolated units (Figure 1C-E), confirming expression of ChR2.

**Figure 1.**
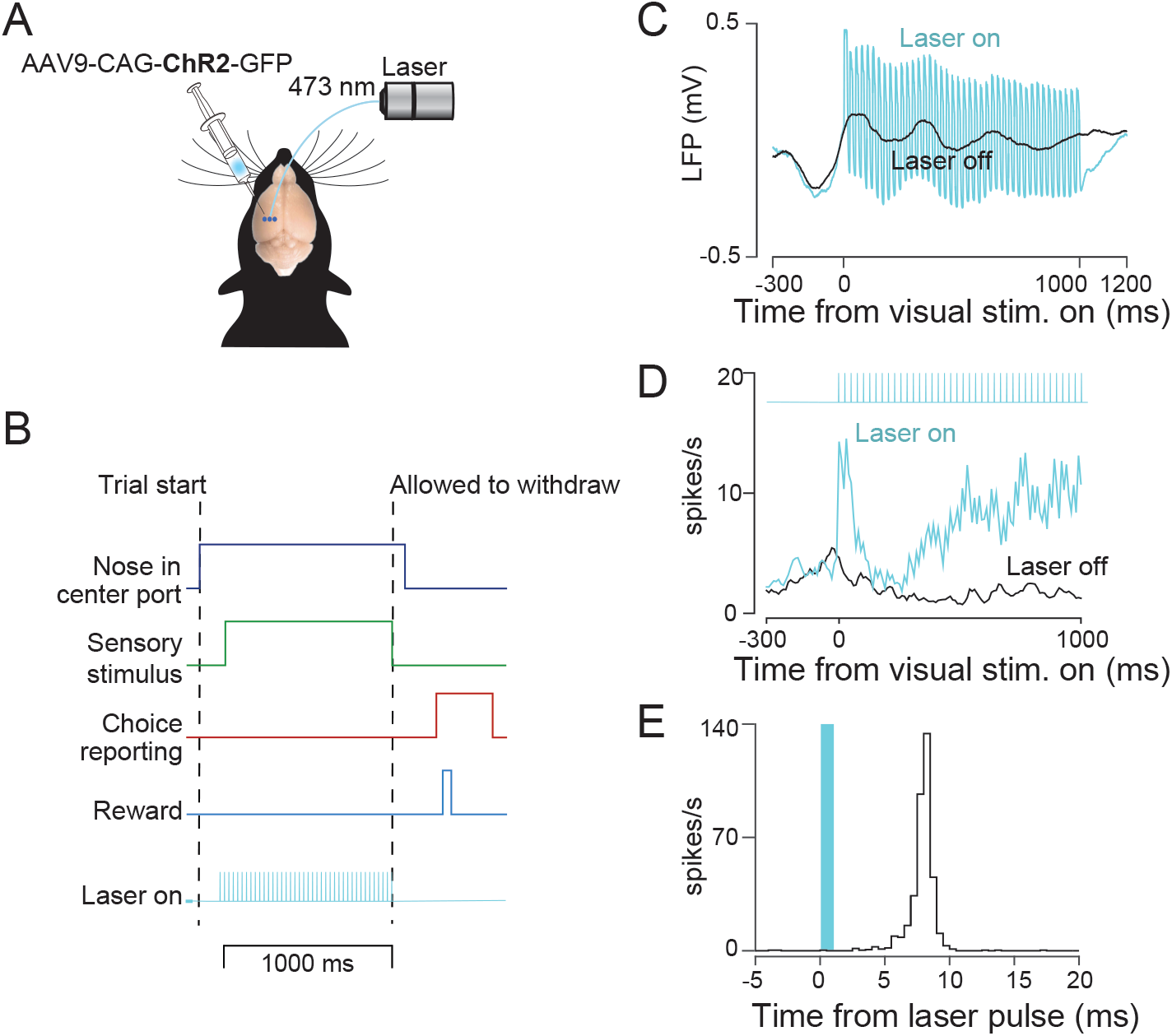
Decision making task and strategy for disrupting PPC activity. (**A**) Schematic of optogenetic approach showing unilateral injections of AAV9-CAG-ChR2-GFP into PPC. (**B**) Schematic of decision-making task. Rats initiated trials by inserting their snouts into a nose poke spanned by an infrared beam (dark blue trace). After a variable delay, a series of auditory or visual events began (green trace). Animals were required to remain in a center port for 1000 ms during which these sensory stimuli were presented. Animals were then allowed to withdraw. They reported choices (red trace) at either a left or right decision port and were rewarded with a drop of water (light blue trace) when correct. Optogenetic stimuli (42Hz, 5-20mW, cyan trace) were presented throughout the 1000 ms period on randomly selected trials. (**C**) LFP recorded on laser-on and laser-off trials via a tetrode attached to the stimulating optical fiber. (**D**) Peristimulus time histogram for an example well-isolated single neuron for laser-on (cyan) and laser-off (black) trials. (**E**) Perievent time histogram in which responses are aligned to individual pulses of blue light.

Stimulation reduced the animal’s decision accuracy, a reduction that was large and significant on visual trials (Figure 2A, left, p=0.0002) and more modest and insignificant on auditory trials (Figure 2A, right, p=0.08). The larger effect on visual decisions was not due to a difference in baseline performance. Animals’ accuracy on control trials was similar for auditory vs. visual decisions: averaged across all stimulus rates, the proportion of correct choices was 0.68 correct for auditory trials vs. 0.70 for visual trials, a difference which did not reach significance (p=0.08, paired t-test). The effects of stimulation were restricted to decision accuracy and did not affect the animals’ ability to engage in the task or to report choices. The proportion of trials that were aborted because of an early withdrawal from the center port did not change appreciably with stimulation and in fact sometimes decreased, indicating improved task engagement (Supp. Table 1). Further, the time it took for animals to withdraw from the center port once the stimulus had ended was similar on stimulation and control trials (response time, Figure 2B and Supp. Table 1). Finally, the time that elapsed between when animals left the center port and when they arrived at a reward port was similar on stimulation and control trials (movement duration, Figure 2C and Supp. Table 1).

**Figure 2.**
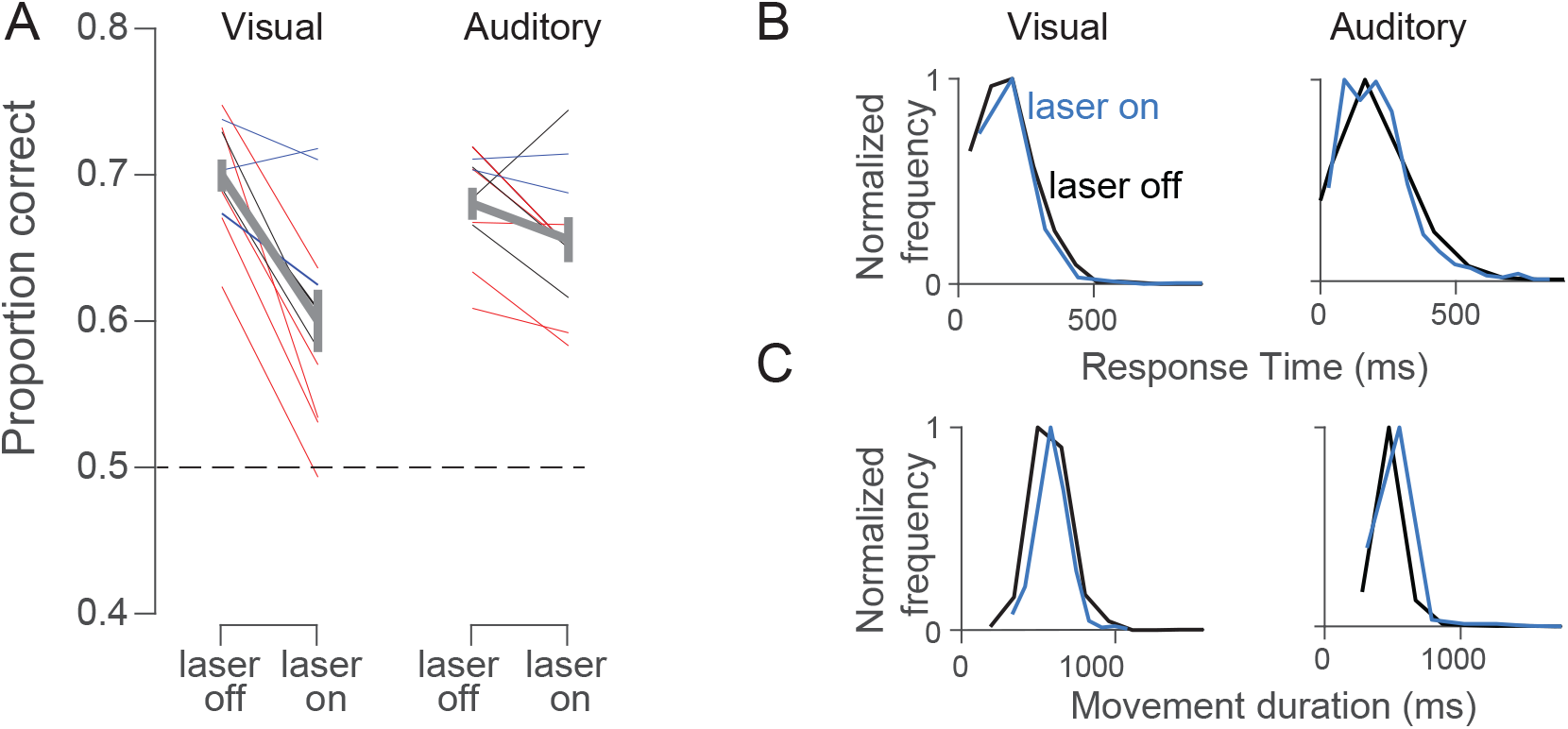
Stimulation drives a strong reduction in visual, but not auditory decisions and largely spares movements. (**A**) Proportion correct for laser on vs. laser off trials for visual (left) and auditory (right) trials. Each line illustrates values for a single site; lines of the same color are from the same animal. Thick gray line indicates mean (±s.e.m.) for all sites. Dashed line indicates chance performance. (**B**) Response times from an example site were similar following laser on (blue) and laser off (black) trials on both visual (left, 183 vs. 184 ms, p=0.16) and auditory (right, 209 vs. 211 ms, p=0.67) decisions. (**C**) Movement durations from an example site were similar following laser on (blue) and laser off (black) trials on both visual (left, 588 vs. 578 ms, p=0.15 ms) and auditory (right, 491 vs. 476 ms, p=0.01) decisions.

The reduction in visual accuracy we observed could be due to disruption of any of a number steps in the process by which the animal converts incoming sensory signals into a decision. To gain insight into which steps in the decision process were disrupted, we visualized the stimulation and control data as psychometric functions, in which the proportion of correct choices is plotted as a function of the stimulus rate (Figure 3A). For visual trials at this example site, the psychometric function on stimulation trials (blue) is shallower than that for control trials (black). This means that a given change in visual stimulus rate (horizontal axis) had a weaker effect on contralateral decisions (vertical axis) on stimulation vs. control trials, a change that contributed to the overall reduction in decision accuracy evident in Figure 2A (left).

**Figure 3.**
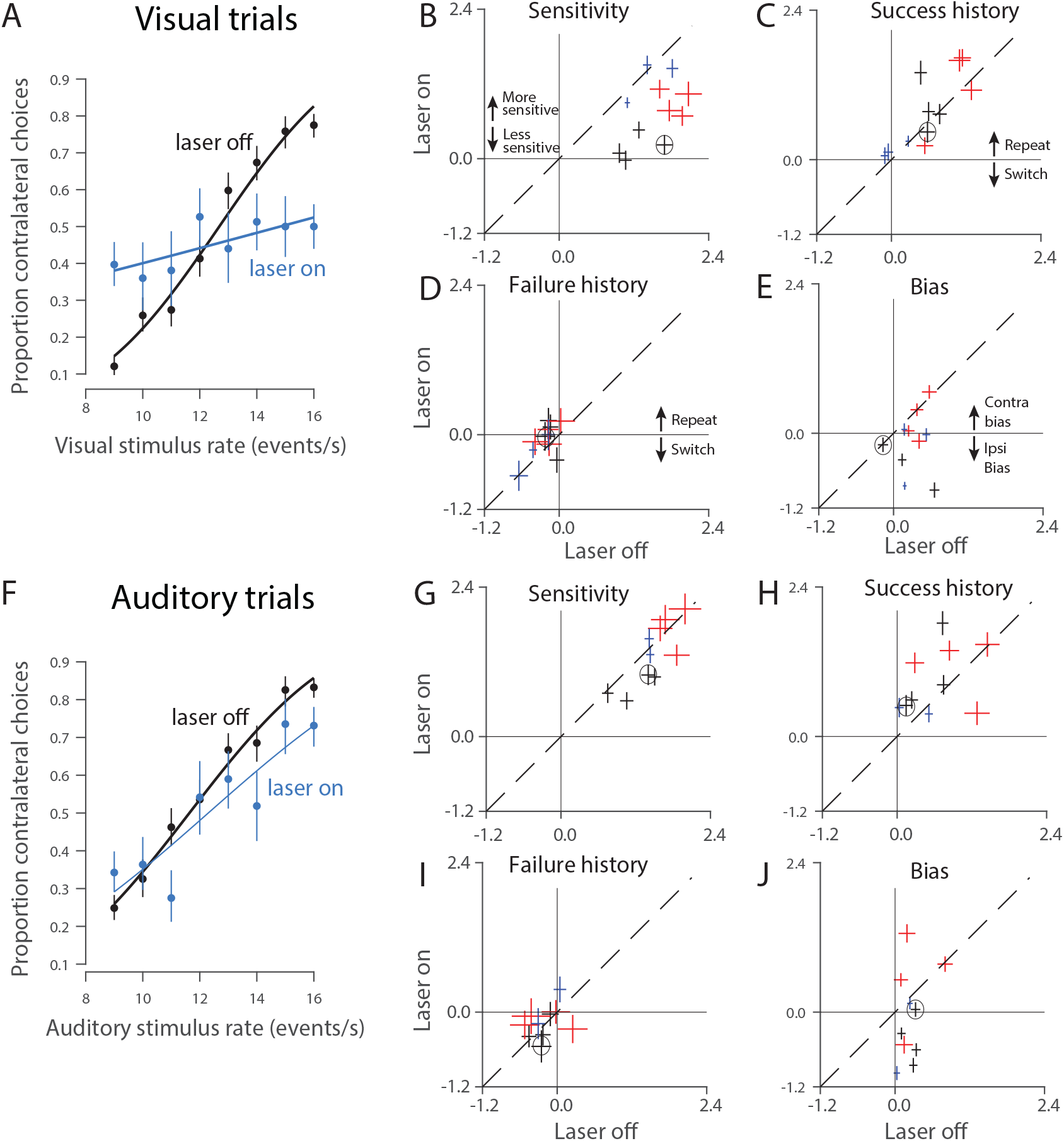
PPC disruption has a larger effect on visual, compared to auditory, decisions. (**A**) Visual psychometric functions from stimulation in a single location within PPC (Rat 1, 1,624 trials). Smooth lines are fits to the data (logistic regression). Error bars reflect the Wilson binomial confidence interval. (**B**) Outcome of a probabilistic model that measures the effect of sensitivity to stimulus rate on decisions. The fitted parameter is plotted for stimulation (laser on, vertical axis) vs. control (laser off, horizontal axis) trials. All values are positive indicating that increasing stimulus rate led to more high rate decisions. Error bars: standard errors. Dashed line, *y* = *x*. Colors: individual rats; multiple points for each animal indicate data collected from different optical fibers/depths (sites) within PPC. Black circle indicates the animal shown in (A). (**C**) Same as (B) but for the “success history” parameter. Positive values indicate the rat tended to repeat rewarded decisions. (**D**) Same as (B) but for the “failure history” parameter. Negative values indicate the rat tended to switch after unrewarded decisions. (**E**) Same as (B) but for the “bias” parameter. Zero indicates unbiased decisions; negative values indicate an ipsilateral bias. (**F**) Auditory psychometric functions from the same rat/site as in (A) (1,655 trials). (**G-J**) Same as (B-E) but for auditory trials.

To quantify and more deeply understand these changes, and to compare them across animals and stimulation sites, we used a probabilistic decision model (Lau and Glimcher, 2005; Busse et al., 2011) (Methods). This model included sensitivity to stimulus rate, bias, and two additional factors that affect decisions (related to trial history, see below). Each factor could in principle be affected by stimulation, offering insight into the precise nature of the observed deficits. The most consistent factor affected by stimulation was reduced sensitivity, evident in all animals and significant overall (Figure 3B, most points below y=x line; p=0.0003; effects were individually significant in 7 of 11 sites; p<0.01, t-test). This loss of sensitivity serves to reduce the steepness of the psychometric function described above (Figure 3A). To test whether the optogenetic stimulation was temporally localized, we examined the sensitivity parameter from stimulation trials (just as above) alongside that from control trials that immediately followed a stimulation trial (Figure Supplement 2). We saw no effect on sensitivity for control trials following stimulation trials (p=0.49), suggesting that the effect of optogenetic stimulation was temporally precise, in keeping with the fast offset we observed in the LFP (Figure 1C). The restriction of sensitivity effects to the current trial confirms that our optogenetic strategy was successful in driving temporally precise disruption.

In addition to a loss of sensitivity, the shallower psychometric functions (and worse accuracy) on stimulation trials might be explained by an increased tendency for rats to be influenced by the previous trial’s outcome. Because trials are generated independently, any influence of the previous trial, such as repeating a successful decision, is deleterious. The probabilistic decision analysis ruled out this explanation. Stimulation had a very weak effect on the degree to which the current decision was influenced by the previous trial’s success (Figure 3C, p=0.16; 2 of 11 individual sites were significant, p<0.01, t-test) or failure (Figure 3D, p=0.04, t-test, 0 of 11 individual sites were significant, p<0.01, t-test). These results rule out two “strategy” explanations for the stimulation effects, supporting the hypothesis that stimulation drove a loss of visual sensitivity. We were unable to find consistent effects of trial history even when we examined the effects of previous left and right decisions separately (Figure Supplement 3).

A final effect of stimulation on visual trials was on the animal’s bias. Bias is defined as a tendency for animals to favor one side over the other regardless of the strength of the sensory evidence. Under the hypothesis that PPC in one hemisphere is preferentially involved in computations relevant to the contralateral side (Crowne et al., 1986; Hanks et al., 2006) disrupting PPC in one hemisphere should bias the animal away from contralateral choices, driving an ipsilateral bias. We observed this ipsilateral bias at a number of sites (Figure 3E, most points below y=x line; p=0.013; effects were individually significant in 6 of 11 sites; p<0.01, t-test).

Altogether, the probabilistic choice analysis suggests that the reduced decision accuracy on visual trials was largely driven by a reduced sensitivity to visual inputs, sometimes exacerbated by a bias away from contralateral choices.

To ensure that the effects observed were due to ChR2 activation, we repeated the same stimulation protocol in a rat not injected with ChR2 (Figure Supplement 4A-C). Similar values were observed on stimulation and control trials for bias (p=0.22, t-test) and sensitivity (p=0.20, t-test). This indicates that blue light in the brain does not by itself drive the effects we observed.

### Optogenetic disruption of PPC spared auditory decision-making

We evaluated performance on interleaved auditory trials to determine whether the effects reported so far reflected vision-specific sensory deficits, or instead reflected more general decision-making deficits. Auditory decisions from the same site and sessions as in Figure 3A demonstrate a much weaker effect of stimulation (Figure 3F). Some sites (4/11) did have small reductions in sensitivity that reached significance (p<0.01; Figure 3G, points below dashed line). Across sites, however, this reduction in sensitivity was not significant (Fig. 3G, p=0.13). Further, a site-by-site comparison revealed that visual sensitivity was significantly more reduced by stimulation compared to auditory sensitivity (Figure Supplement 5A; p=0.0021, t-test).

No consistent effect was observed on animals’ reliance on trial history, whether it was a previous trial’s success (Figure 3H, p=0.16, t-test) or failure (Figure 3I, p=0.28, t-test). The effect on bias was idiosyncratic. As with visual trials, an ipsilateral bias was sometimes present, but biases in the opposite direction were also observed (Figure 3J; a significant ipsilateral bias (same direction as for visual trial) was evident at 6 of 10 sites and a significant contralateral bias was evident at 2 of 10 individual sites, p<0.01, t-test). No significant change was present overall (Figure 3J, p=0.13, t-test). The difference in bias between auditory and visual trials did not reach significance (Figure Supplement 5B, p= 0.29, t-test).

### PPC neurons are more strongly driven by individual visual events than by auditory events

The consistent effect of stimulation on visual, but not auditory, sensitivity is evidence against a simple model in which auditory and visual signals equally influence PPC (Figure 4A). Our results suggest a new class of model in which PPC is a key player for translating visual, but not auditory, sensory signals into decisions (Figure 4B,C). To provide an independent test of this class of model, we evaluated whether individual visual sensory events had a larger impact on PPC responses compared to individual auditory sensory events using a previously collected, large scale (N= 101,972 successful trials) electrophysiological dataset (Raposo et al., 2014). This sensory-evoked response is potentially separate from the decision-related responses reported in previous analyses (Raposo et al., 2014), which focused on slower signals evolving over an entire 1000 ms decision (Figure 5A) rather than on transient responses following individual sensory events. Indeed, a signature of individual events can be obscured when trials with events at different times are averaged, especially when the slower decision modulation is large (as in Figure 5A), or with wide-filter smoothing (as is often used to improve signal-to-noise ratios). We evaluated the impact of individual auditory and visual events by aligning electrophysiological responses to individual visual or auditory events in single neurons and removing the slow component (Methods). Many neurons were driven by individual stimulus events (Figure 5B). This event modulation was frequently evident in visual trials (84 of 317 neurons at p<0.01), but only occasionally evident in auditory trials (5 of 317 n eurons at p<0.01). Modulation was significantly more common due to visual compared to auditory events (p≪10^−4^, χ^2^ 2×2 contingency table). Modulation was also significantly stronger for visual compared to auditory events within neurons (Figure 5C,D, p≪10^−4^, Figure supplement 6, paired sign test). Importantly, a larger effect of visual inputs was evident despite the fact that auditory and visual stimuli were carefully matched so that they had an equivalent effect on decisions (See figure 1c of Raposo et al., 2014). These observations, like the disruption effects, support models in which visual inputs are dominant in PPC (Figure 4B,C).

**Figure 4.**
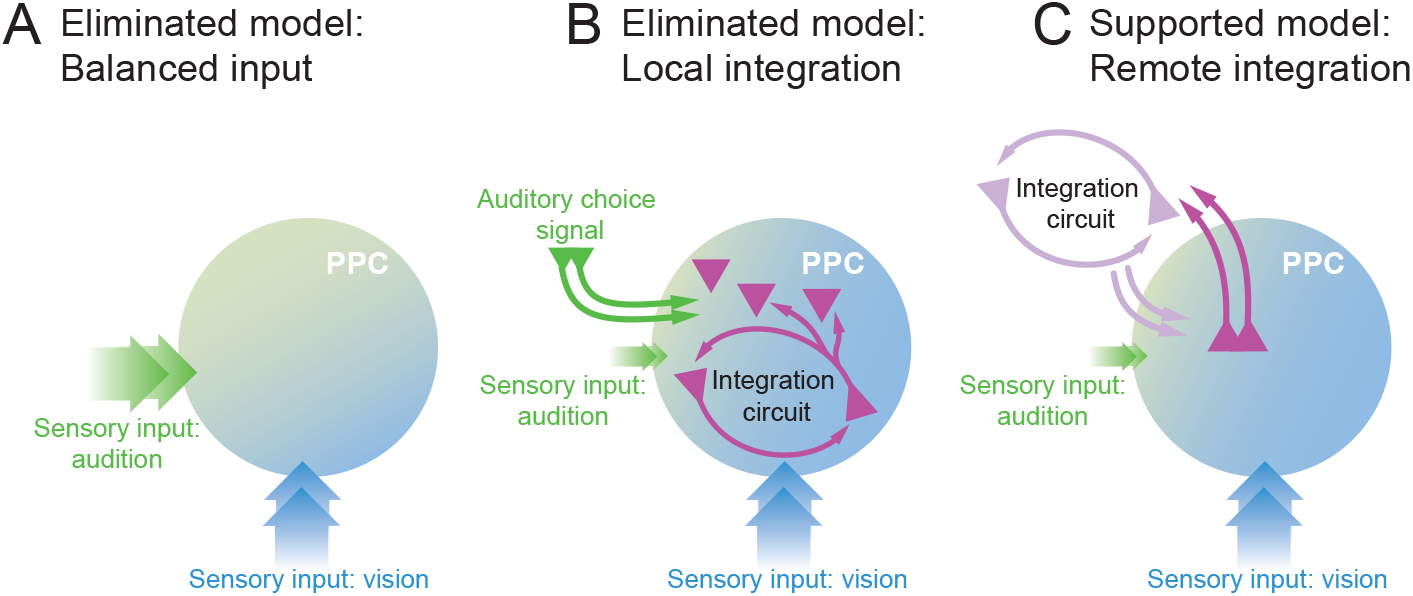
Putative models for PPC’s role in a decision circuit. (**A**) Balanced input model eliminated by the disruption experiment. (**B**) Local integration model in which visual inputs to PPC are stronger than auditory inputs and evidence over time is integrated within PPC. (**C**) Remote integration model in which visual inputs to PPC are stronger than auditory inputs and evidence over time is integrated at a remote location and fed back to PPC.

**Figure 5.**
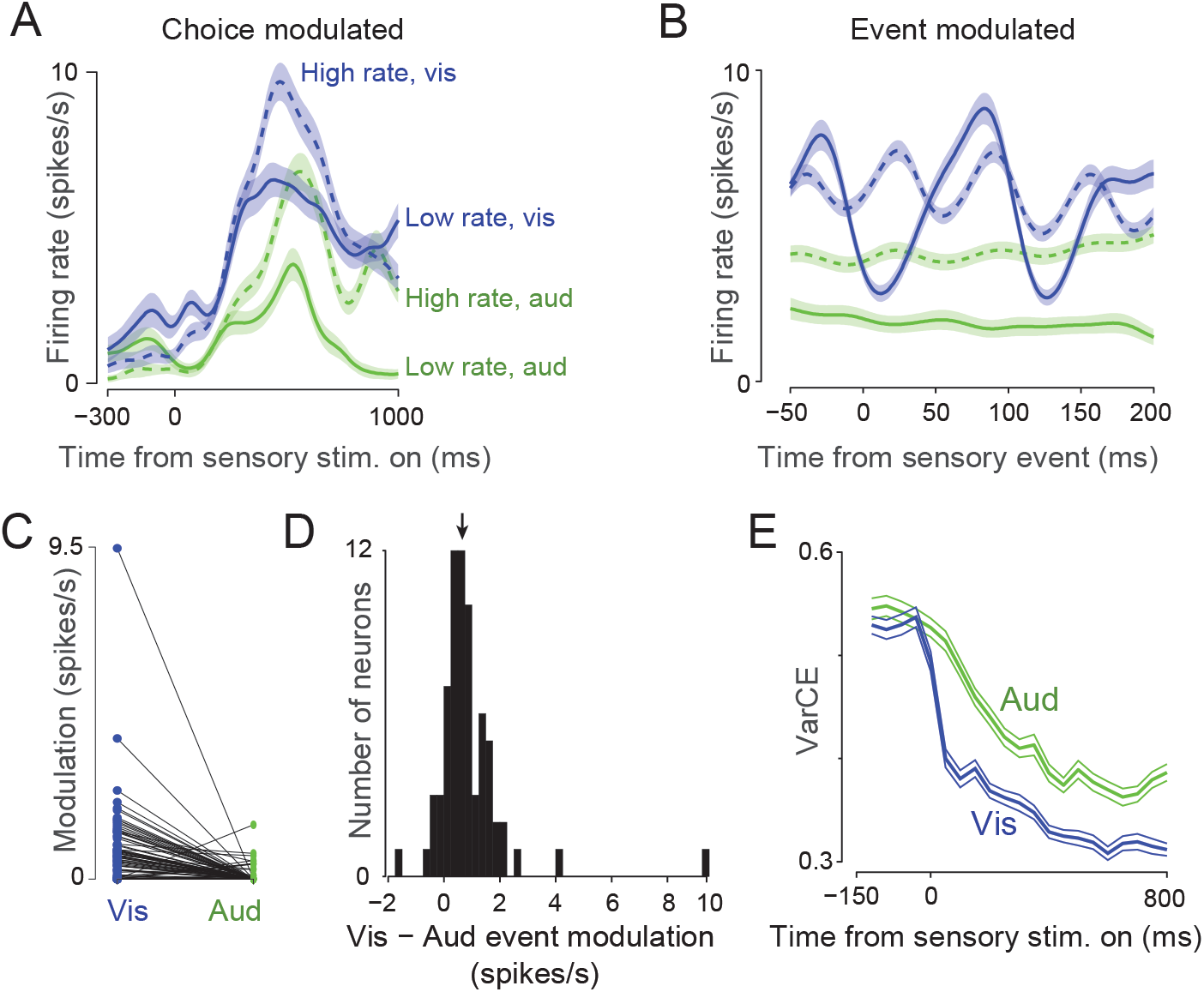
Electrophysiological analyses suggest PPC discriminates individual visual events and does not act as an evidence accumulator. (**A**) Trial-averaged peristimulus time histogram for an example neuron. Solid traces, low-rate trials; dashed traces, high-rate trials; blue traces, visual trials; green traces, auditory trials. Transparent fills show s.e.m. The outcome of the decision (dashed vs. solid lines) and the stimulus modality (blue vs. green lines) drove slow modulations over the 1000 ms decision. (**B**) Peri-event histogram for same neuron as in (A), aligned to individual visual or auditory events (Methods). Same conventions as (A). (**C**) Modulation strength of each neuron by visual (blue) and auditory (green) events. Example dataset shown for low-rate trials only. The two measurements for each neuron are connected by a line. Note that after correcting for noise, many neurons had a modulation index of 0. (**D**) Histogram over neurons of the modulation index for visual minus the index for auditory. Same dataset as (C). Neurons with both modulation indices equal to 0 were excluded. Arrow shows median (0.68; p<10^−10^, sign test). (**E**) VarCE computed relative to stimulus onset for auditory (green) and visual (blue) trials.

### Analysis of trial-to-trial variability suggests sensory signals are accumulated remotely

Visual inputs to PPC might be integrated locally within the area (Local integration model; Figure 4B), or might be integrated in another region which projects back to PPC (Remote integration model; Figure 4C). Theory suggests that as integrators accumulate evidence, they should also accumulate noise that will vary across trials (Churchland et al., 2011). Thus, trial-to-trial variability should increase during decision formation in areas that reflect evidence accumulation. By contrast, in areas that reflect sensory inputs or movement preparation, the stabilizing influence of the input or plan will drive decreases in trial-to-trial variability over time in a trial (Churchland et al., 2010; Rajan et al., 2010). A measure of trial-to-trial variability in neural responses, the Variance of the Conditional Expectation (VarCE, Churchland et al., 2011) was designed to distinguish these possibilities. In several decision-related brain areas in the monkey, the VarCE has been demonstrated to increase over time during evidence accumulation decisions (Churchland et al., 2011; Ding, 2015). In the present data, clear decreases in VarCE were observed for both auditory and visual trials (Figure 5E).

## Discussion

Our results argue that PPC discriminates visual inputs which are accumulated off-site to guide decision-making. Three observations support this. First, optogenetic disruption of PPC reduces sensitivity on visual decisions but largely spares auditory decisions and does not affect movement metrics or behavioral strategies. Second, individual visual events drive larger electrophysiological responses in single neurons compared to auditory events, even though these events are equally effective in driving behavior. Finally, trial-to-trial variability decreases during decisions, suggesting the presence of a stabilizing influence from sensory inputs or action planning rather than the destabilizing influence of evidence accumulation. Taken together, these findings point to PPC as required to discriminate visual signals which are then accumulated remotely (Figure 4C).

The experimental design here allowed us to go far beyond previous disruption studies because we used temporally precise disruption, included multiple sensory modalities, and analyzed the decisions with a probabilistic choice model. This experimental design allows us to understand the nature of the deficit with precision, and suggests that the rats were deficient in discriminating visual stimuli. This deficit could have arisen because of a number of changes in early visual processing circuits, including reduced signal-to-noise or a reduction in attention. A role in attention and prioritizing space have long been attributed to PPC in primates (For reviews, see Colby and Goldberg, 1999; Gottlieb, 2007). Having established that PPC plays a causal role in visual decisions, future studies can aim to uncover the computations performed in PPC that support these decisions.

The evidence-accumulation signals apparent in electrophysiological recordings (Hanks et al., 2015) may reflect feedback from other areas (Figure 4C). This offers an explanation for why inactivation during auditory decisions has little effect (Erlich et al., 2015) despite strong modulation of PPC neurons during such decisions (Hanks et al., 2015). This feedback possibility is further supported here by our measure of trial-to-trial variability, the VarCE, which is diagnostic of underlying neural computations. VarCE increases in areas that reflect accumulation of evidence, such as primate lateral intraparietal area (Churchland et al., 2011), caudate nucleus and frontal eye field (Ding, 2015). By contrast, VarCE decreases in areas that reflect sensory input or motion planning (Churchland et al., 2010), since those computations push the network towards a more stabilized state (Rajan et al., 2010). The decreasing VarCE observed here may likewise indicate stabilization. Future studies will be needed to determine whether this stabilization is driven by sensory input, action planning or both. We speculate that the sharper decrease in the VarCE seen on visual decisions (Figure 5E, blue) may reflect the dual stabilizing influences of visual sensory input and action planning feedback, while the slower and less deep decrease of the VarCE on auditory decisions (Figure 5E, green) may reflect only a single stabilizing influence, most likely action planning feedback. The idea that PPC neurons reflect, in part, action planning signals is in keeping with previous observations that the direction and magnitude of decision-related tuning does not depend strongly on whether decisions were instructed by auditory or visual inputs (Raposo et al., 2014).

Disrupting neural activity to determine a structure’s role in behavior, as we have done here, can lead to challenges in interpretation (Otchy et al., 2015). Fortunately, a number of aspects of our experimental design bolstered our ability to interpret these disruption experiments. First, we minimized the chance that the rats would detect the disruption and adjust their strategy: the disruption was transient and present only on a minority of randomly selected trials. This allowed us to rule out that the sparing of auditory decisions was explained by reliance on alternate circuits during slow timecourse inactivation, a possibility left open by other studies of decisions (Raposo et al., 2014; Erlich et al., 2015; Katz et al., 2016). Second, we disrupted activity by artificially elevating firing rates, a method that is ideal for disruption of behaviors that depend on heterogeneous and time-varying population codes (Churchland and Shenoy, 2007; Roberts et al., 2012; Rodgers and DeWeese, 2014; Otchy et al., 2015). For such behaviors, optogenetic stimulation offers advantages over optogenetic suppression because it introduces a new, aberrant signal. This may more strongly perturb the population code compared to suppression, especially because the overall change to the population can be larger than for suppression (which suffers floor effects).

A final aspect of our experimental design that aids interpretation of effects is that we studied decisions guided by two different sensory modalities. This allowed us to rule out some alternatives to the possibility that PPC disruption reduces visual sensitivity. For instance, one alternative explanation for the deficits during visual decisions is that PPC stimulation activated neurons in downstream areas, disrupting circuits that plan the actions needed for decision reporting. We can rule out these “off target” effects (Otchy et al., 2015) because auditory decisions, which would rely on the same motor circuits, were largely spared. However, one off-target effect we cannot fully rule out is on primary visual cortex. If PPC has denser feedback projections to primary visual cortex than to primary auditory cortex, PPC stimulation might have stronger off-target effects on primary visual cortex neurons, explaining the largely visual deficits we observed. Fortunately, an independent support for a role of PPC in discriminating visual events is provided by our observation from electrophysiology that visual inputs more strongly drive the temporally precise PPC responses that are needed to discriminate visual inputs.

An additional caveat is that the extent of the disruption due to direct activation is not known with absolute precision. This is because although we measured neural activity during stimulation, the spatial coverage of our electrodes was insufficient to determine at what distance from the stimulating electrode the blue light ceased to activate neurons. Fortunately, for optogenetic disruption, the spatial extent of activation is primarily determined by parameters of the stimulation: wavelength, fiber diameter, numerical aperture, and laser power. This is unlike chemogenetic inactivation in which the spatial extent of activation depends on the spread of viral infection; this is also unlike pharmacological disruption, in which the spatial extent of activation depends on diffusion of the reagent. To estimate light spread, and thus the spatial extent of our disruption, we used published calculators (Methods). The measurements made to estimate light spread using these calculators are extensive, including measurements both in vivo (Guo et al., 2014) and in slice (Aravanis et al., 2007; Huber et al., 2008).

One possibility that these calculations deem unlikely is that the blue light (and thus the direct activation) spread to primary visual cortex (V1). Our stimulating fiber was positioned at 3.8 mm posterior to Bregma. According to the calculations from the Brain Light Tissue Transmitter, the irradiance at 1.15 mm away from the fiber is expected to be 0.5 mW/mm^2^, too weak to drive neurons (Guo et al., 2014). This distance would be ~4.9 mm posterior to Bregma, at which the very most anterior tip of V1 is 1.5 mm lateral to where we positioned our optetrode (Paxinos and Watson, 2007). Even if a small number of V1 neurons were somehow affected, our full-field stimulus would only have altered the response of a few V1 neurons representing the extreme lower nasal edge of one hemifield. Activation of V1 neurons is therefore very unlikely to be responsible for our behavioral effects.

Although we think it unlikely that our results were due to direct stimulation of V1 neurons, it is essential to acknowledge that outstanding questions remain in understanding the relationship between PPC, classically defined by its thalamic inputs (Chandler et al., 1992; Reep et al., 1994), and the secondary visual areas that are observed via anatomical tracing (Coogan and Burkhalter, 1990; Montero, 1993). The shallower psychometric functions we observed on stimulation trials (Figure 3A) are reminiscent of those seen during inactivation of extrastriate regions in monkey (See Figure 2c of Katz et al., 2016). One possibility is that rat PPC shares features with monkey extrastriate regions, such as a causal role in processing raw visual inputs (Newsome and Pare, 1988; Katz et al., 2016). Alternatively, rat PPC may be akin to monkey PPC (Brody and Hanks, 2016), and the extrastriate-like deficits we observed are present because the PPC coordinates used by us and others (Whitlock et al., 2012; Raposo et al., 2014; Erlich et al., 2015) encompass separate, more extrastriate-like areas. Challenges in distinguishing a candidate structure from its nearby neighbors have long been acknowledged. The present results make clear that at least some of this cortical territory is causally involved in visual decision making. However, improved resolution of areas and their borders using methods such as widefield retinotopic mapping (Schuett et al., 2002; Andermann et al., 2011; Garrett et al., 2014; Glickfeld et al., 2014) and noise analyses (Kiani et al., 2015) combined with high-density recordings may inform further experiments narrowing down the key areas for decision making in cortex.

In conclusion, we demonstrate that PPC plays a causal role specifically in visual decision-making. Our results are in keeping with previous inactivation studies, but allowed us to more deeply probe the effects of inactivation by ruling out alternative explanations for the deficits to visual decision-making. Further, our analysis of electrophysiological responses provides independent evidence of a dominant role for vision in PPC. By establishing PPC as part of a circuit for visual decision-making, we pave the way for future studies that will reveal how visual signals within PPC are transformed as they are passed to subsequent areas.

## Materials and Methods

### Animal subjects

All experimental procedures were in accordance with the National Institutes of Health’s *Guide for the Care and Use of Laboratory Animals* and approved by the Cold Spring Harbor Animal Care and Use Committee. Adult male Long Evans rats (200-250g, Taconic Farms) were housed with free access to food and restricted access to water starting from the onset of behavioral training. Rats were housed on a standard (nonreversed) light dark cycle; experiments were run during the light part of the cycle. Rats were pair-housed initially, but were singly housed once they received injections or implants (below).

### Behavior

Four rats were trained on a rate discrimination task (Figure 1B) described previously (Raposo et al., 2012; Sheppard et al., 2013). Briefly, rats were trained to judge whether the overall rate of a repeating auditory (click) or visual (flash) stimulus was high or low compared to a learned category boundary (12.5 events/second). Three of the four rats were trained that rightwards choices were rewarded following high rate stimuli and leftwards choices were rewarded following low rate stimuli; one rat was trained with the opposite contingency. Stimuli were presented over 1000 ms during which time the rats had to remain in a central port. After this time, rats indicated their choice on each trial by moving to a left or right reward port. Response time (Figure 2B) is the time between when the stimulus ended and when the rat departed the port. Movement time (Figure 2C) is the time between exiting the center port and entering a reward port. Movements to the correct reward port yielded a drop of water (10-25μL).

Animal training typically began 2-3 weeks following viral injection of ChR2; the training period lasted 5-6 weeks and was completed before implanting the stimulation/recording assembly (see *Viral Injection* and *Implants for Electrophysiology*, below).

### General surgical procedures

All rats subject to surgery were anesthetized with isoflurane and administered 5 mg / kg ketoprofen before surgery for analgesia. Isoflurane anesthesia was maintained by monitoring respiration and foot pinch responses throughout the surgical procedure. Ophthalmic ointment was applied to keep the eyes moistened throughout surgery. Lidocaine solution (~0.1 mL) was injected below the scalp to provide local analgesia prior to performing scalp incisions. 0.05 mg / kg buprenorphine was administered daily for post-surgery analgesia (usually 2-3 days).

### Viral injection

We induced ChR2 expression in the left PPC of 3 rats using adeno-associated virus (AAV, serotype 9) carrying the gene ChR2 fused with green fluorescent protein (GFP) under the control of the CAG promoter (AAV9-CAG-ChR2-GFP). This promoter induces the expression of ChR2 in all cell types. Unilateral injections of this construct were made in the left PPC of 4-6 week old rats. We made 2-3 separate penetrations along the medial-lateral axis with the goal of maximizing expression in PPC and minimizing the spread outside of this area. Stereotactic coordinates (relative to Bregma) for Rats 1 and 2 were −3.8 mm AP, −2.2 /−3.2 /−4.2 mm ML; and for Rat 3 were −3.8 mm AP, 2.2/3.7 mm ML. We made a small craniotomy and positioned a calibrated glass pipette within the craniotomy, perpendicular to the brain’s surface. For the injection, we applied pressure to a syringe that was attached to the pipette via plastic tubing. Injections were made at 400, 600 and 800 μm below the pial surface. At each depth, 140 nL was injected. We refrained from deeper injections to avoid viral spread to subcortical structures. Histology obtained at the end of the experiment (Figure Supplement 1) indicated robust virus expression.

Note that because all injections were in the left hemisphere, lateralized effects are referred to as “ipsi” or “contra” because these mean the same for all animals. Because we trained different animals to associate the left vs. right port with low rate choices, all behavioral data is plotted relative to the injected hemisphere. This convention makes it possible to distinguish biases towards a particular side from biases towards a particular rate because “high rate” trials are not always associated with the same side.

### Implants for electrophysiology

Rats were implanted with custom optetrode (Anikeeva et al., 2012) implants that were prepared in-house. Each assembly contained up to 8 independently moveable tetrodes (Nickel/chrome alloy wire, 12.7 μm, Sandvik–Kanthal). Tetrodes were connected to an EIB-36 narrow connector board (Neuralynx) mounted on the implant assembly. Six to eight of the tetrodes were attached to optical fibers used for delivering light (Anikeeva et al., 2012; Znamenskiy and Zador, 2013). Optical fibers were 62.5 μm in diameter with a 50-μm core. In 2 of the rats, fiber tips were sharpened to a point with a diamond wheel to improve tissue penetration and increase the angular spread of the light exit cone. Each optical fiber was glued to a tetrode; the pair was mounted on an independently moveable microdrive. The assembly was secured within a plastic enclosure prior to implanting. Tetrodes were gold-plated to 300-700 kΩ at 1 kHz; one additional tetrode was used as an internal reference for electrophysiological recordings and plated to ~100 kΩ.

For implantation during surgery, we followed procedures described previously (Raposo et al., 2014). Briefly, we positioned the entire optetrode assembly so as to center it relative to the previously made injections (−3.8 mm AP and 2.5 mm ML). A durotomy was performed and the implant assembly was lowered until the tetrodes just penetrated the pial surface. 2% agarose solution was applied to cover the tetrodes and craniotomy, and dental acrylic (Lang Dental) was applied to secure the implant to the skull. The incision was closed around the base of the implant using 1-2 Vicryl sutures anterior and posterior to the implant. Following surgery, tetrodes were advanced in increments of 40-80 μm until action potentials were encountered.

### Optogenetic stimulation

We used blue light (473 nm) with intensity ranging from 5-20mW at the fiber tip. To estimate the spread of light, we used a well-established method, the <u>Brain Light Tissue Transmitter</u>, which estimates light spread based on wavelength, fiber diameter, numerical aperture and power. We elected to use this method for three reasons. First, the estimates of light spread are accurate and reliable because the systematic way in which the measurements were collected to generate the calculator (multiple measurements at each of many distances from fiber tip). Second, the estimates from the calculator are in accordance with many additional published measurements for blue light spread in rodents, both in vivo (Guo et al., 2014) and in slice (Aravanis et al., 2007; Huber et al., 2008). The measurements in slice afford a very precise estimate because experimenters can directly measure light spread by placing the slice over the photodetector of a power meter. Finally, the use of published measurements is justified because the spread of light is likely to be homogeneous across animals; light spread mainly depends on the properties of brain tissue. As a result, judging the extent of stimulation is more straightforward compared to judging the spread of chemogenetic disruption. In chemogenetic disruption (Rogan and Roth, 2011), all infected neurons are activated with equal probability by the ligand to a synthetic receptor delivered by a virus. Therefore the extent of stimulation is determined mainly by the extent of viral spread, so quantifying the spread is essential. Indeed, we estimated spread of effect in just this way in our previous paper (Raposo et al., 2014). In the current study, extent of expression is less informative because neurons expressing ChR2 that are beyond the range of the blue light will be unaffected. Even if we had expressed ChR2 non-specifically across the brain, we still would have achieved specificity because of the natural restriction of the blue light. Indeed, studies routinely achieve specificity by performing optogenetic stimulation in animals expressing ChR2 brain-wide (e.g. Guo et al., 2014).

Based on the diameter of the fiber (50μm) and its numerical aperture (0.22), we estimate (http://web.stanford.edu/group/dlab/optogenetics/, 2015) that at a distance of 0.5 mm away, irradiance was 24.8 mW/mm^2^ and at a distance of 1.15 mm away from the fiber, irradiance was below 0.5 mW/mm^2^. Given that 0.5 mW/mm^2^ has been shown to be the minimal required intensity to induce spiking intensity in awake animals (Guo et al., 2014), we infer that our stimulation mainly affected ChR2 expressing neurons within this range. In all animals, we confirmed that stimulation elicited a clear change in the LFP on at least one tetrode (Figure 1C) although typically, responses were observed on multiple tetrodes.

On a subset of randomly selected trials (“stimulation trials”, 15–25%) we delivered blue light to activate ChR2-expressing neurons in PPC, using a 473 nm diode-pumped solid-state (DPSS) laser. On these trials the laser was triggered at the beginning of the stimulus presentation (visual or auditory) and was kept on throughout the entire decision formation period (1000 ms), delivering light pulses at a rate of 42 Hz (Figure 1D). On the remaining trials (“control trials”, 75–85%) no optical stimulation occurred. We used two techniques to minimize the rats’ ability to detect the optical trials by seeing the blue light. In Rats 1 and 2, we covered the implant with black insulating tape before beginning each session. To ensure that no light was emitted, the experimenter would deliver light into the laser while it was connected to the animal in a dark booth and visually inspect the implant for any escaping light. A second method was developed because adding and removing tape from the implant daily reduced the integrity of the implant and sometimes resulted in premature explantation. In this second method, the implant was not covered in tape, but we also used a second optical fiber in a ferrule *not* implanted in the brain (that is, light from this laser was blocked from entering the brain). Light from this second fiber was still visible and thus served to mask the light from the stimulation. The second fiber was illuminated on every trial (control trials and stimulation trials). As a result, the presence of blue light would be difficult to use to detect optical stimulation trials.

We typically collected data from a single “site” (stimulation on one fiber at a particular depth) for 5-8 days; behavioral data were pooled over those days. We then either stimulated on a different tetrode or advanced the current optetrode at least 200μm.

### Analysis of stimulation effects

We measured the effects of stimulation on multiple aspects of behavior. First, to systematically determine the effects of stimulation on four factors contributing to the animal’s decision, we used a probabilistic decision model (Busse et al., 2011):

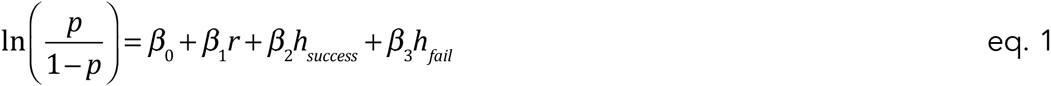

where *p* is the probability of making a rightward decision, *r* is the stimulus strength (its rate relative to the 12.5 Hz category boundary; the true range of −3.5 to 3.5 events/s above and below the boundary were scaled so that values ranged from −1 to 1), *h_success_* indicates whether the previous trial was a success (1 if the right side was rewarded, −1 if the left side was rewarded; 0 otherwise) and *h_fail_* indicates whether the previous trial was a failure (1 if the failure followed a decision to the right side, −1 if the failure followed a decision to the left side; 0 otherwise). The coefficients were fit in Matlab (Mathworks, Natick MA) using glmfit and a logit linking function. The observer’s decision was predicted as a combination of four factors, so the values of the fitted coefficients (*β*_0–3_) provide insight into how much each parameter of the model influences the decision on any given trial. Stimulation (laser-on) and control (laser-off) trials were fitted separately so that the coefficients could be compared. In principle, only the rate (*r*) should influence the rat’s decision because this determines the reward contingency, but previous work has shown that in practice, side bias and reward history bias are influential. If animals were to rely more on reward history bias on stimulation trials, this would have reduced their overall performance since the correct response for the current trial is independent of the previous trial. Therefore, this analysis afforded a deeper insight into the factors reducing the rat’s accuracy on stimulation trials.

To assess significance of differences in the fitted coefficients (Figure 3B-E,G-J; Figure supplement 4B,C), two tests were performed. First, we conducted one-sided paired t-tests for each site separately. The effect of stimulation was evaluated for each of the four fitted parameters: bias, sensitivity, success history and failure history. The t-statistic for each was computed directly using the values of the fitted parameters and their associated standard error returned by glmfit (computed from the square root of the diagonal values of the covariance matrix). The standard error on the difference was calculated by propagating the error associated with the stimulation and no-stimulation values of each parameter. Second, we conducted one-sided paired t-tests for the data pooled across all sites and animals.

We also measured the effect of stimulation on other behavioral measures. First, we determined whether stimulation changed the animal’s ability to remain in the center port for the required 1000 ms duration. We used a χ^2^ test to evaluate whether the proportion of trials in which the animal withdrew early differed for stimulation versus control trials (Supp. Table 1). Second, we evaluated the time that elapsed between when the stimulus ended and when the animal exited the center port (response time). We used an unpaired, one-sided t-test to evaluate whether response times differed for stimulation and control trials (Figure 2B, left, Supp. Table 1). Finally, we evaluated the time that elapsed between when the rat left the center port and when it arrived at one of the two side ports (movement duration). We used an unpaired, one-sided t-test to evaluate whether movement duration differed for stimulation and control trials (Figure 2C, right, Supp. Table 1).

We performed two additional analyses of electrophysiological responses from a previously collected dataset (Raposo et al., 2014). We used these instead of the electrophysiological dataset associated with optogenetic stimulation because of its large size (n=101,972 trials). Although we did record well-isolated neurons with the optetrodes used here (Figure 1C), the population size did not provide the statistical power needed for the relevant analyses. Because the previous dataset was trained on rats performing an identical task, the data was ideally suited to these analyses.

### 1. Analysis of the effect of single stimulus events on neural responses

For the first analysis (Figure 5B-D), we wished to determine whether single flash or click events modulated firing rates at fast timescales independent of overall condition modulation (the tuning captured in a typical peri-stimulus time histogram). To do so, we considered all successful trials of the highest or lowest rates, separately. We first smoothed each trial’s firing rate with an acausal Gaussian (15 ms SD). We then made peri-event time histograms (PETHs) for low-rate trials. The first three events of every trial were discarded to reduce the effects of onset transients and adaptation. To remove the slower tuning component, we performed a linear detrending of this PETH using 1.5 cycles of the stimulus. Use of 1.5 cycles helped reduce slope bias when fitting a periodic waveform. Next, as a raw estimate of modulation (*m_raw_*), we computed the standard deviation of this waveform over one cycle. However, this estimate is biased: noisier PETHs will tend to produce higher SDs. To correct for this, we assumed that the observed spiking was the sum of the “true” underlying fluctuations in the firing rate plus a noise process (Shadlen and Newsome, 1998; Nawrot et al., 2008; Churchland et al., 2011). Conveniently, the variance of the noise process at any time point is simply the square of the s.e.m (denoted 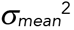). Correcting for the noise exactly is a challenging problem in Cox process inference, but since event modulation was typically small compared to total firing rate, we could easily obtain a reasonable approximation of the total noise by averaging the s.e.m. over time points. Since variances add for the sum of independent random processes, our modulation index was therefore:

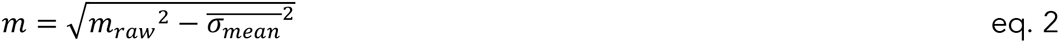

For neurons where 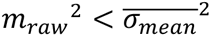, we assigned *m* = 0. To obtain the modulation index for high-rate trials, the same process was repeated using events from high-rate trials. Note that values of exactly 0 were excluded from the histograms in Figure 5D and Figure Supplement 6, and thus the histograms for high-rate trials (which elicited much less modulation) contain far fewer points.

To assess significance for each neuron, we wished to know how often we underestimated the noise process such that the measured modulation was actually no larger than the noise. Since our modulation measure is based on the standard deviation, the relevant comparison is with the standard error of the standard deviation of the noise process. For a Gaussian random process, this is: 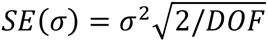, where *DOF* is the number of degrees of freedom and here 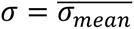. Since our trace was smoothed, *DOF* ≠ *N*−1. Instead, we used a common estimator for *DOF* of a smoothed series: Tr(*S_λ_*), where *S_λ_* is the linear smoothing matrix and Tr(·) indicates taking the trace of a matrix. Using this approximation, for a Gaussian smoothing kernel,

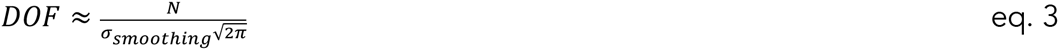

To obtain a p-value for each neuron, we performed a one-tailed Z-test of the neuron’s modulation index against the Gaussian distribution with mean 0 and standard deviation 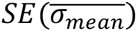 for that neuron. By inspection of individual neurons, event modulation was almost always greater for low-rate trials, so only low-rate trials were tested for significance. These p-values were not corrected for multiple comparisons because the goal was to obtain an estimate of the number of modulated neurons, not to determine whether any neurons were modulated. We also tried using a Bonferroni-corrected p-value threshold for the latter purpose: at a significance level of p<0.05/317, 45 neurons were significant for visual and 2 for auditory.

### 2. Analysis of trial-to-trial variance to provide insight on neural computation

To understand how trial-to-trial variability evolved over the course of auditory and visual decisions, we computed a measure of spike count variability, the variance of the conditional expectation (VarCE, Churchland et al., 2011; Brostek et al., 2013; Marcos et al., 2013; Ding, 2015). Briefly, this measure assumes that the total measured spike count variance can be divided into 2 components using the law of total variance for doubly stochastic processes: (1) variance of counts that would be produced by a stochastic point process with a given rate (“spiking noise”), and (2) the variance of the rates that would produce those counts (“conditional expectation”). The VarCE isolates the second of these components and is therefore informative about underlying mechanism. In principle, the VarCE is computed by subtracting an estimate of the first component from the total spike count variance:

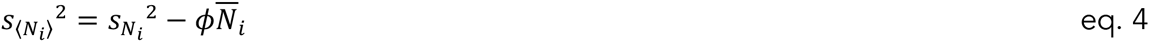

where *N_i_* is a vector of spike counts for a given neuron and given condition in time window *i, s_N_i__*^2^ is the sample variance of those spike counts, 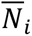 is the mean spike count of a neuron across trials of a given condition in time window *i*, and *ϕ* is a constant that approximates the degree to which spike count variability scales with firing rate (Geisler and Albrecht, 1995; Nawrot et al., 2008). In practice, as in previous work, we computed *ϕ* separately for each neuron in the dataset by taking the minimum of the measured Fano factor across all conditions and time points. To make it possible to combine data from multiple conditions, we estimated *s_N_i__*^2^ using the residuals — that is, by subtracting from each sample count the mean for all trials sharing its condition. The VarCE plotted in Figure 5e is the variance of the union of residuals from all conditions, minus the weighted average of the stochastic variance 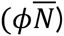(see eq. 6, Churchland et al., 2011). A sliding time window with a width of 100 ms was used for the traces in Figure 5E. Longer (150 ms) and shorter (60 ms) windows yielded similar results.

## Author contributions

All authors designed the experiments, A.L., M.R. and J.S. carried out the experiments, D.R., A.C. and M.K. designed the analyses, A.C. wrote the paper.

## Acknowledgements

We thank Barry Burbach and Simon Musall for advice on the figures and analyses and Bernardo Sabatini for providing comments on a draft of the paper. Funding for this work was provided by NIH EY022979 (AKC), the McKnight Foundation (AKC), the Simons Collaboration on the Global Brain (AKC, MTK), the Pew Foundation (AKC), the Klingenstein Foundation (AKC) and the Marie Robertson Memorial Fund of Cold Spring Harbor Laboratory (AKC).

